# Wenzhou shrimp virus 8 (WzSV8) detection by unique inclusions in shrimp hepatopancreatic E-cells and by RT-PCR

**DOI:** 10.1101/2022.08.15.504032

**Authors:** Jiraporn Srisala, Dararat Thaiue, Piyachat Saguanrut, Suparat Taengchaiyaphum, Timothy W. Flegel, Kallaya Sritunyalucksana

## Abstract

The genome sequence of Wenzhou shrimp virus 8 (WzSV8) (GenBank record KX883984.1) was described in 2015 from wide screening for RNA viruses in aquatic animals. A closely related sequence (GenBank record OK662577.1) from the whiteleg shrimp *Penaeus vannamei* was deposited in 2021 under the name Penaeus vannamei picornavirus (PvPV). In 2022 another closely related sequence (GenBank accession: OP265432) was submitted under the name Penaeus vannamei solinvivirus (PvSV). In 2021, prior to the publication of PvPV and PvSV, we used an RT-PCR method devised from the sequence of KX883984.1 (described herein) to screen for WzSV8 in specimens of cultivated shrimp. Samples that gave positive RT-PCR results were subsequently tested by *in situ* hybridization (ISH) analysis to identify virus target tissues. Several tissues gave positive ISH results within morphologically normal nuclei. Thus, these tissues were of no use for diagnosis of WzSV8 by normal histological analysis. However, unique basophilic, cytoplasmic inclusions within vacuoles in the hepatopancreatic E-cells were also found in the same WzSV8-positive shrimp specimens, sometimes accompanied by a smaller eosinophilic inclusion. We call these Lightner double Inclusions (LDI) that can be considered pathognomonic for diagnosis of WzSV8 infection when detected using the light microscope. Although no current proof of WzSV8 is the cause of disease, it is important to investigate new viruses and related tissue anomalies, even from normal cultivated shrimp, to determine whether they may have any relationship to significant negative effects on the production of cultivated shrimp.

## INTRODUCTION

Wenzhou shrimp virus 8 (WzSV8) (Li et al. 2015) was discovered in 2015 by wide screening of marine animals for RNA viruses using high throughput sequencing and the resulting GenBank record (KX883984.1) of the full genome sequence is 10,445 nucleotides. A more recent publication from China also gives another full genome sequence (GenBank record OK662577.1) that is highly similar to that of WzSV8 (97% coverage and 95.4% sequence identity), but under the newly proposed name Penaeus vannamei picornavirus (PvPV) with a full genome sequence of 10,550 nucleotides (Liu et al., 2021). The authors placed PvPV in the Order Picornavirales, Family *Dicistroviridae* as a positive-sense, ssRNA virus. In 2022, yet another highly similar sequence (GenBank accession: OP265432) was submitted under the name Penaeus vannamei solinvivirus (PvSV) (Cruz-Flores et al. 2022). We received a copy of this publication only during the review process for this report.

Although the PvPV article contained no histological analysis, it did include an electron micrograph of a cytoplasmic viral inclusion within a vacuole of an unspecified hepatopancreatic epithelial cell (Liu et al., 2021). In contrast, the PvSV article did include histological analysis accompanied by ISH test results, but they differed from those reported herein. Specifically, the ISH positive signals were present only in the nuclei of cells that appeared normal by hematoxylin and eosin (H&E) staining. In contrast to the publication by Cruz-Flores et al. (2022), we found that some WzSV8-positive shrimp specimens also showed, in addition to normal cells with ISH-positive nuclei, other unique histopathological lesions that can be considered pathognomonic for WzSV8 infection. These pathognomonic lesions would be useful for histopathologists who commonly use the light microscope in routine screening for pathogens. In addition, histopathological analysis can be used for screening archived paraffin blocks for microscopic detection of WzSV8 while RT-PCR analysis and ISH might not be an option because of the instability of RNA.

After the publication reporting WzSV8 (Li et al., 2015) and a subsequent publication from Australia reporting the presence of WzSV8 in the transcriptome of wild *Penaeus monodon* (Huerlimann et al., 2018), we used the sequence of KX883984.1 to design an RT-PCR detection method for WzSV8. We then used the RT-PCR protocol with fresh shrimp specimens submitted by clients to screen for the presence of WzSV8. This involved taking samples for RNA extraction together with tissue preparation for subsequent histopathological analysis. Tissue samples from shrimp positive for WzSV8 by RT-PCR were then subjected to ISH analysis to locate WzSV8 infected cells and reveal their morphology. Parallel tissue sections stained with hematoxylin and eosin (H&E) could then be examined to determine whether H&E stained lesions characteristic of WzSV8 could be identified.

Because WzSV8 is an RNA virus, archived shrimp specimens preserved in paraffin blocks for histological analysis cannot be used for ISH analysis because of the instability of RNA. Care must even be taken with a short period of fixation in Davidson’s fixative to obtain reliable ISH results. Thus, in the current absence of a validated immunohistochemical method, long-storage, paraffin-embedded tissues cannot be used for molecular analysis. These limitations do not apply to analysis of the unique lesions by H&E staining.

The purpose of this report is to provide information allowing for the detection of WzSV8 infections using standard light microscope methods for examination of shrimp hepatopancreatic tissue sections stained with hematoxylin and eosin. An RT-PCR protocol to screen for WzSV8 is also provided.

## MATERIALS AND METHODS

### Sample sources and overall research protocol

Samples from global clients from the Americas and the Indo-Pacific were used as RNA templates to develop a nested RT-PCR method for WzSV8. The forward and reverse primers for the first RT-PCR and the nested PCR were designed from the sequences of GenBank record KX883984.1. This was done prior to our knowledge of the publication on *Pv*PV (Liu et al., 2021) and submission of its genome sequence (OK662577.1) to GenBank. The RT-PCR amplicons from three specimens from the Indo-Pacific (SG1.1, SG1.2, and SG 1.3) and from the Americas (SG2.1, SG2.2, and SG2.3) were subjected to sequencing and bioinformatics analysis. The exact source of our specimens is confidential client information. Our clients have been fully informed of our results. If any reporting to national competent authorities is required, it is the client’s responsibility to do so. It is then the responsibility of the respective national competent authority to report, in turn, to any international organization which they may have committed to do so.

### Histological analysis

Standard methods were used for shrimp fixation and processing to prepare hematoxylin and eosin (H&E) stained tissue sections (Bell and Lightner, 1988). These were analyzed using a Leica ICC50 HD digital light microscope. For semi-thin sections, tissue samples were removed from the outer, E-cell region of the hepatopancreas for fixation in 4% glutaraldehyde solution (4% glutaraldehyde, 19 mM NaH_2_PO_4_°H_2_O, 81 mM Na_2_HPO_4_, pH 7.4), embedded in epoxy resin, sectioned using a Leica EM UC6 ultramicrotome with a glass knife and stained with toluidine blue as previously described (Sriurairatana et al., 2014).

### RT-PCR method

The first step RT-PCR reaction is performed in 12.5 μl mixture consisting of 1X Reaction Mix (Invitrogen, USA), 0.4 μM each of WzSV8-482F and WzSV8-482R primers (Table 1), 0.5 μl of SuperScript III RT/Platinum Taq Mix (Invitrogen, USA) and 100 ng of RNA template. The RT-PCR protocol begins with 50°C for 30 min followed by 94°C for 2 min and then by 35 cycles of 94°C for 30 sec, 60°C for 30 sec and 68°C for 45 sec plus a final extension at 68°C for 5 min. For the nested PCR step, the 12.5 μl mixture contains 1X OneTaq Hot Start Master Mix (NEB, USA), 0.2 μM of each WzSV8-168F and WzSV8-168R primer (**Table 1**), and l μl of the product solution from the first RT-PCR step. The nested PCR protocol is 94°C for 5 min, followed by 25 cycles of 94ºC for 30 sec, 60ºC for 30 sec and 72ºC for 30 sec plus a final extension for 5 min at 72 ºC. The amplicons yielded are 482 bp and 168 bp, respectively.

**Table 1.**
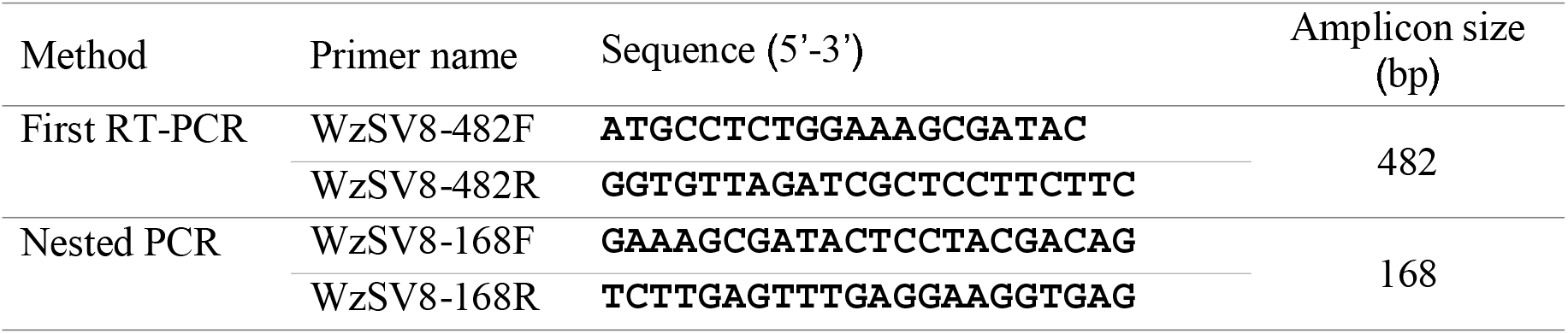
Primers used for the nested RT-PCR detection method for WzSV8 developed in this study.

### Bioinformatics analysis

Multiple sequence alignment of amplicons SG1.1, SG1.2, SG1.3, SG2.1, SG2.2, and SG2.3 was performed by Clustal Omega (https://www.ebi.ac.uk/Tools/msa/clustal/), and nucleotide sequence similarity was analyzed using the BLASTn sequence analysis tool (https://blast.ncbi.nlm.nih.gov/Blast.cgi). The phylogenetic tree was constructed using MEGA 7 program. Tree topology was evaluated using bootstrap analysis by the maximum likelihood method with default parameters for 1000 replicates.

### *In situ* hybridization (ISH)

ISH assays were carried out as previously described (Srisala et al., 2021). Briefly, the primers WzSV8-482F and WzSV8-482R (Table 1) were used with a plasmid template containing cDNA of a WzSV8 genome fragment to prepare a DIG-labeled, DNA probe for WzSV8. The negative controls consisted of adjacent tissue sections that were treated the same as the test samples, except for omission of the DIG-labeled probe. Each ISH included 3 adjacent tissue sections, one for H&E staining, one for the ISH probe and one for a no-probe control.

## RESULTS AND DISCUSSION

### Nested RT-PCR detection of WzSV8

The primers designed for WzSV8 were based on the nucleotide sequence record of GenBank accession no. KX883984.1. A schematic diagram representing primer regions specific to a portion upstream (nt 3889-nt 4370) of the putative RNA-dependent RNA polymerase (RdRp) is shown in **Fig. 1**. The sensitivity of the method is 20 copies per reaction vial using purified WzSV8 amplicons as the template. An agarose gel showing RT-PCR amplicons obtained from 3 samples each from the Indo-Pacific (SG1.1, SG1.2, and SG 1.3) and the Americas (SG2.1, SG2.2, and SG2.3) is shown in **Fig. 2**.

**Figure 1.**
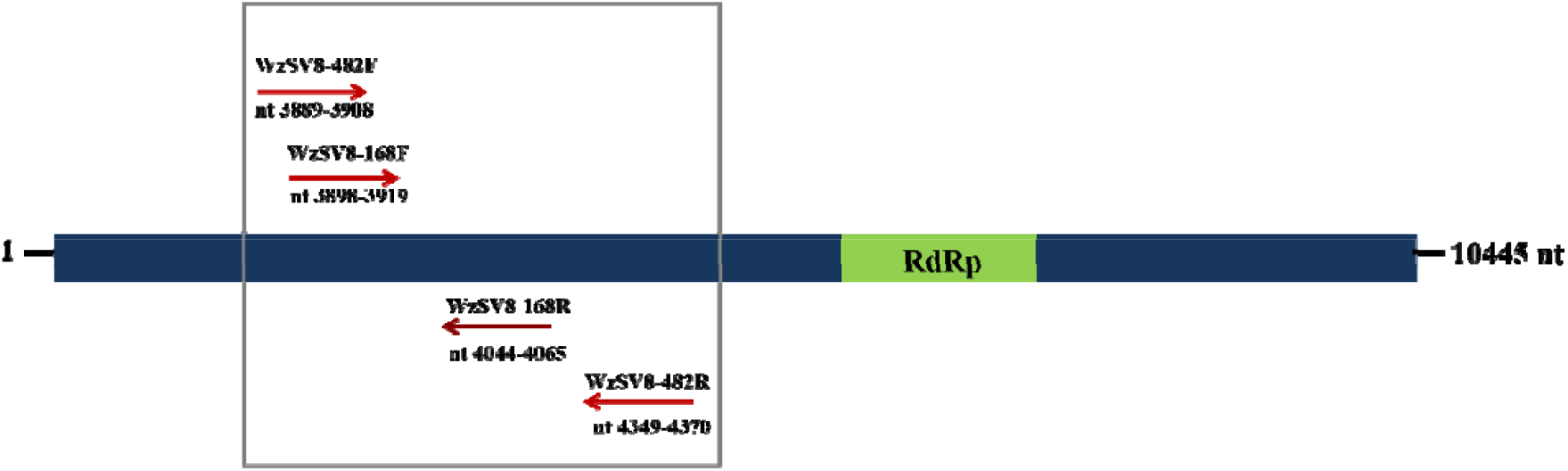
A schematic diagram representing primer regions upstream of RdRP gene The red arrows indicate the locations of the primers for the first RT-PCR step WzSV8-482F and WzSV8-482R, and the nested PCR step WzSV8-168F and WzSV8-168R The nucleotide positions of the primers according to the position of the GenBank record KX883984 1 are shown

**Figure 2.**
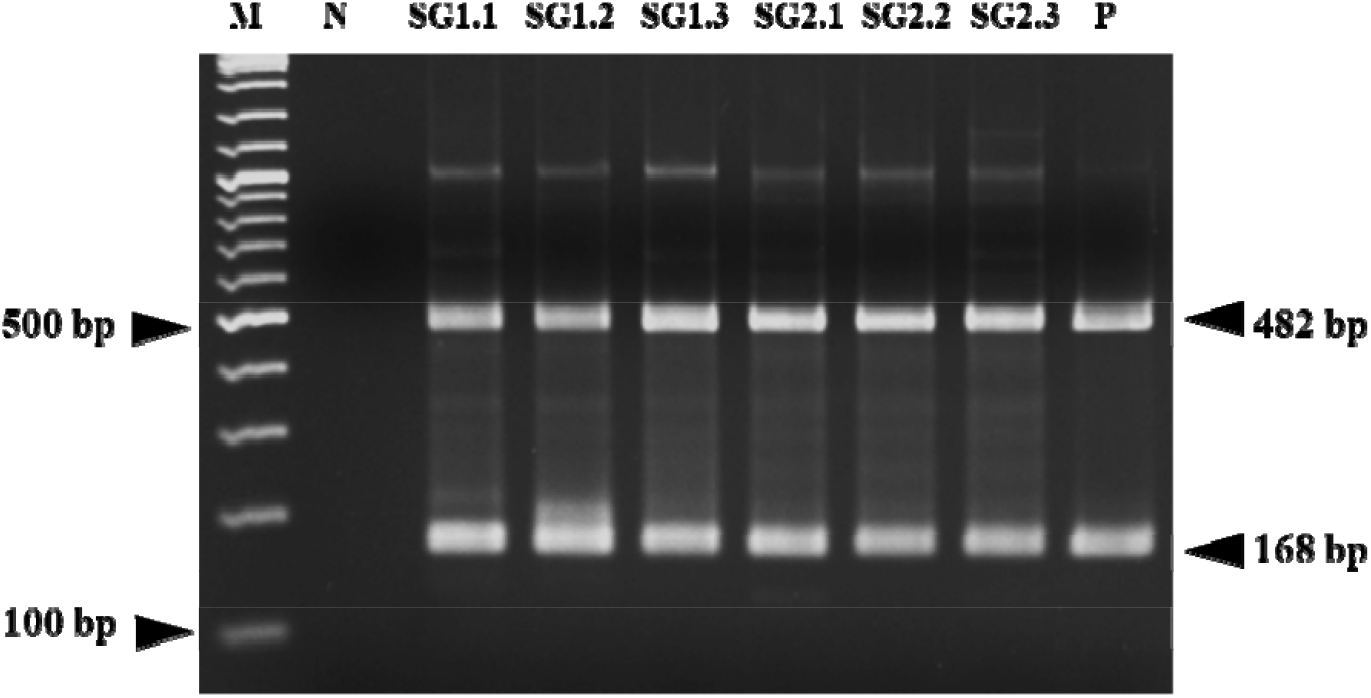
Photograph of an agarose gel showing RT-PCR amplicons obtained from 6 shrimp specimens using the WzSV8 method M 2 log DNA marker; N Negative control; P Positive control

A Clustal Omega alignment of the consensus sequences of these amplicons with the matching regions of the WzSV8 sequence (GenBank record KX883984.1) and the PvPV sequence GenBank record OK662577 1 is shown in **Supplementary Fig S1** The identity among the eight nucleotide sequences differed appreciably **Table 2**, even for the two GenBank sequences from China 93 3-98 1 This indicates relatively high variability in only 6 different WzSV8 samples that may possibly differ in virulence for shrimp Phylogenetic analysis revealed that the WzSV8 sequences of the Indo-Pacific specimens SG1 fell in a clade with the Chinese sequences and separated from the American sequences SG2 **Fig 3**

**Table 2.** Percent identity matrix among the eight aligned nucleotide sequences created by Clustal Omega program.

|  | SG1.1 | SG1.2 | SG1.3 | SG2.1 | SG2.2 | SG2.3 | OK662 | KX883 |
| --- | --- | --- | --- | --- | --- | --- | --- | --- |
| SG1.1 | 100.0 |  |  |  |  |  |  |  |
| SG1.2 | 92.9 | 100.0 |  |  |  |  |  |  |
| SG1.3 | 93.1 | 99.7 | 100.0 |  |  |  |  |  |
| SG2.1 | 93.9 | 92.9 | 93.1 | 100.0 |  |  |  |  |
| SG2.2 | 93.7 | 93.5 | 93.7 | 99.3 | 100.0 |  |  |  |
| SG2.3 | 92.9 | 91.7 | 91.9 | 97.7 | 97.5 | 100.0 |  |  |
| OK662577.1 | 95.6 | 94.4 | 94.6 | 93.9 | 94.6 | 93.3 | 100.0 |  |
| KX883984.1 | 96.6 | 95.6 | 95.8 | 94.8 | 95.4 | 94.1 | 98.1 | 100.0 |

**Figure 3.**
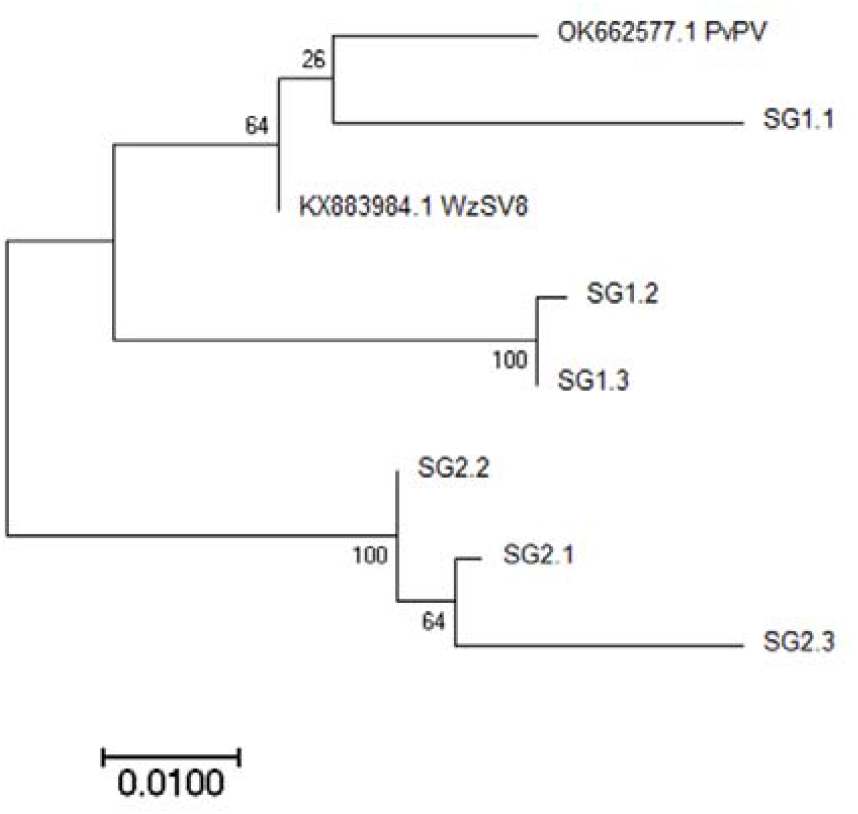
Phylogenetic tree based on RT PCR amplicon sequences of 6 shrimp specimens compared to matching regions of two WzSV8 nucleotide sequences at GenBank. The tree was constructed using the maximum likelihood method with MEGA 7 software. The percentages of replicate trees in which the associated taxa clustered together in the bootstrap test (1000 replicates) are shown next to the branches.

Since our RT-PCR method was developed based on the GenBank record KX883984.1, it may not be suitable for detection of all types of WzSV8 that exist, but we found it useful for specimens from Asia to the Americas and it may serve as a preliminary method until a more universal method can be developed, perhaps to cover all variants. Upon request to the corresponding author, we are willing to provide a free plasmid containing the target for our method. It can be used to transform *E. coli* as a perpetual source of a positive control plasmid for PCR tests and for use as a template to prepare ISH probes. With proper acknowledgement of the source, the plasmid may be distributed freely without seeking prior permission. We wish this tool to be distributed as widely and quickly as possible to encourage cooperation and exchange of information on WzSV8-like viruses.

-

### Identification of WzSV8 lesions in shrimp tissues by ISH

Our first set of shrimp positive for WzSV8 by RT-PCR were processed for histological analysis by H&E staining and by ISH analysis using a DIG-labeled probe prepared by PCR using the plasmid containing the WzSV8 target sequence. Prior to our ISH tests, we did not know the target tissue for WzSV8 except for one electron micrograph in the paper on PvPV (Liu et al. 2021) showing a viral inclusion in a vacuole of an unidentified HP cell. Examples of positive ISH reactions for WzSV8 in an HP sample are shown (black arrows) in a low magnification photomicrograph (**Fig. 4**). This gives a positional reference for the higher magnification photomicrographs that follow. The figure also shows the lack of ISH reactions in the no-probe control. The section shown in Fig. 4 is a tangential section of the rounded, outer region of the HP. The tubules of the HP extend outward in a radial manner from the central (proximal) region of the HP to its outer (distal) perimeter. Thus, the area at the center of this HP section (indicated by an asterisk in Fig. 4) is deeper in the HP than the edges of the image. The center is characterized by the presence of prominent vacuoles of differentiated B-cells and R-cells. In the area around the asterisk, the tubules have been cut perpendicular to the tubule axis (i.e., cross-cut).

**Figure 4.**
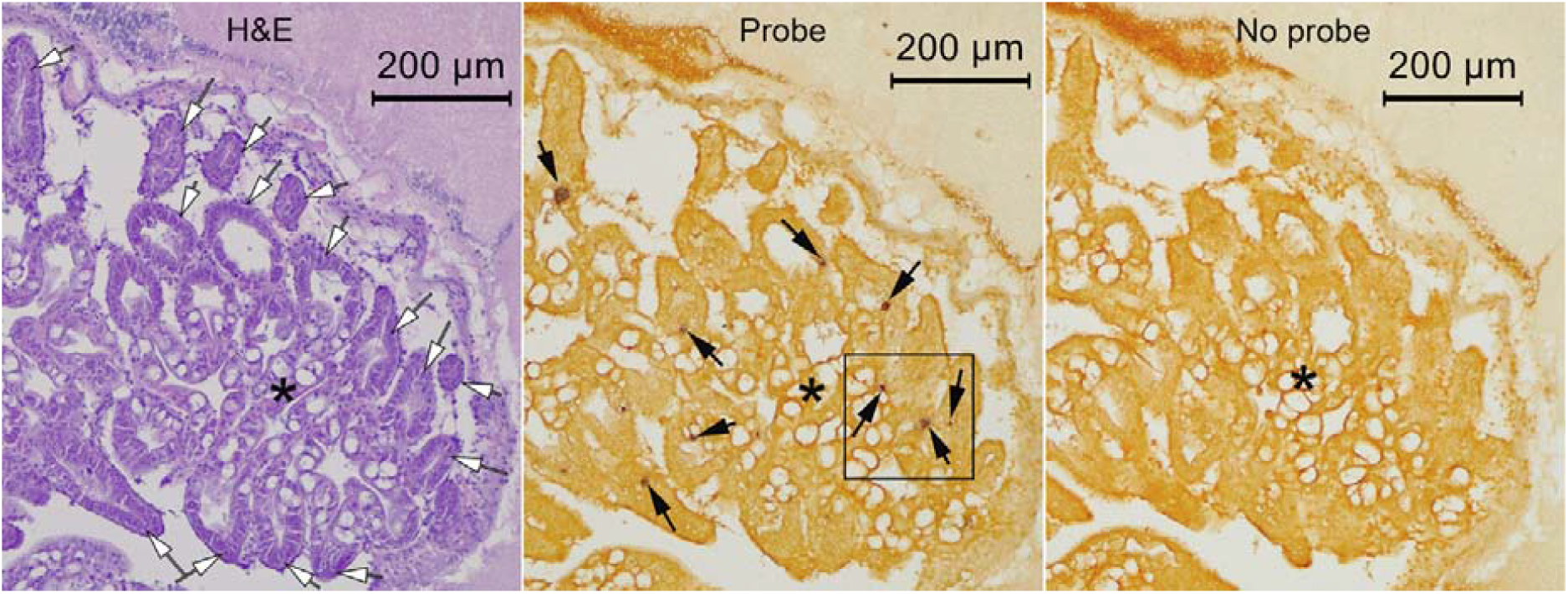
Low magnification photomicrographs from an ISH assay, showing positive ISH reactions (black arrows). These are often found predominantly in E-cells and less frequently in areas (asterisks) of differentiated HP tubule epithelial cells characterized by the presence of vacuoles. The box in the middle of photomicrograph is shown at higher magnification in Fig. 5.

In contrast, moving radially outward from the central area of the section, the tubules are cut more tangential to their long axis and look finger-like, terminating in rounded, conical ends (white-filled arrows) that show cells predominantly without vacuoles. Herein, we refer to these densely stained areas with no to low vacuoles as “E-cell” regions of the HP. The tubule tips themselves include only E-cells that are normally characterized by lack of vacuoles and by the presence of mitotic spindles. Moving from these distal tubule tips towards the proximal end of the tubule, the E-cells begin to differentiate into B, F and R cells and mitotic spindles are not seen.

In the samples we examined, we found that the positive ISH reactions were predominantly in the distal area of the HP i e, in or near the E-cell region rather than in the more proximal differentiated region However, this varied greatly from specimen to specimen A portion of the ISH photomicrograph in Fig 4 box outline is shown at higher magnification in **Fig 5**, where it can be seen that 2 of the positive ISH signals in the E-cells arise from circular inclusions within vacuoles which do not normally occur in E-cells Also in Fig 5 is an E-cell with a positive ISH reaction in both the nucleus and cytoplasm of a cell that has otherwise normal morphology and would not be recognizable as an abnormal cell by H&E staining Additional ISH positive circular inclusions within vacuoles in the cytoplasm of E-cells are shown in **Fig 6** collected from other samples

**Figure 5.**
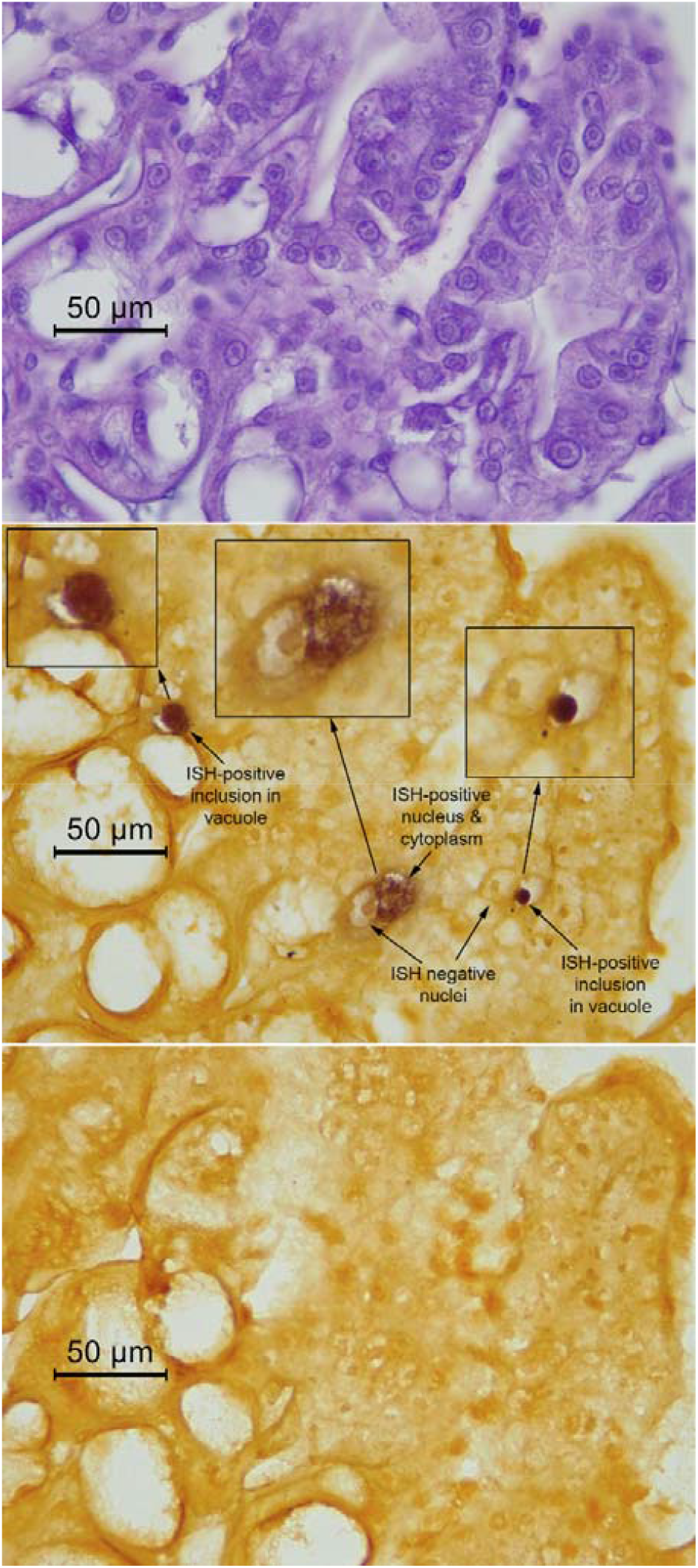
Magnification of the box in Fig 4 showing 2 positive ISH reactions as dark, circular positive signals in vacuoles, resembling the position and shape of basophilic inclusions seen in vacuoles with H&E staining Also shown is another ISH positive cell with no circular inclusion but instead positive ISH reactions in both the nucleus and cytoplasm This positive cell would not differ from other normal cells with H&E staining

**Figure 6.**
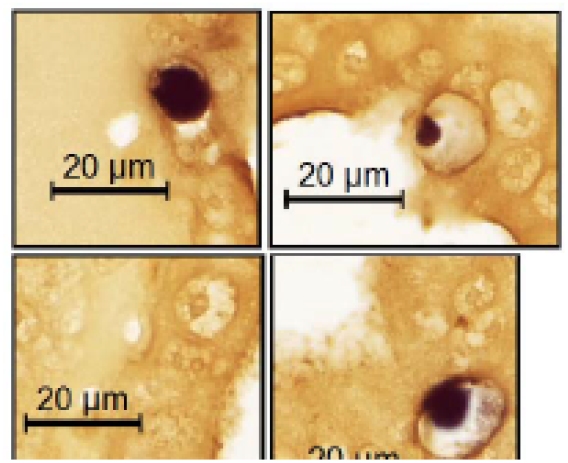
A collection of 4 photomicrographs from various samples showing high magnifications of ISH positive reactions with circular inclusions within vacuoles of E-cells. The sizes of the inclusions range from approximately 3 to 11 μM.

Although we could not find three adjacent tissue sections that showed the same cell with the same inclusion in all 3, we were able to establish a consistent and unique morphological pattern for at least some of the WzSV8 lesions. This consisted of a deeply basophilic, circular inclusion, most distinctively within a clearly defined cytoplasmic vacuole of tubule epithelial cells in the E-cell region of the shrimp HP where vacuoles normally do not occur. Although these lesions also occurred in tubule epithelial cells in the medial and proximal regions of the HP, the presence of basophilic cytoplasmic, vacuolar inclusions in the E-cell region have not previously been reported for shrimp, so they provide a convenient focus for rapid histological detection of WzSV8 infections.

Some of the specimens RT-PCR positive for WzSV8 were grossly normal and obtained from normal harvest ponds. Others, including tissue sections on archived slides or new slides prepared from archived paraffin blocks, originated from diseased shrimp. For example, some exhibited pathological lesions caused by bacteria and/or the microsporidian *Enterocytozoon hepatopenaei* (EHP) in the medial and proximal areas of the HP. Sometimes, the lymphoid organs showed the presence of spheroids. However, these lesions gave negative ISH reactions with the WzSV8 probe (not shown).

### Some specimens gave positive ISH reactions for WzSV8 in normal nuclei

In some of our specimens positive for WzSV8 by RT-PCR, there were occasionally ISH positive signals in nuclei of normal tubule epithelial cells in the medial and proximal areas of the HP (**Supplementary Fig. 2**) and in the subcuticular epithelium, and especially the subcuticular epithelium of the stomach (**Supplementary Figs. S3 & S4)**. Such ISH reactions rarely occurred in the E-cell region. In the adjacent sections stained with H&E in S2-S4, the nuclei and the cytoplasm are histologically normal (i.e., no visible cytopathic effects or lesions). This type of ISH reaction within morphologically normal nuclei was found to occur in some specimens in which the abnormal cytoplasmic, circular inclusions shown in Figs. 4 to 8, 10 and 11 were absent. However, both types of ISH reactions occurred together in other specimens. The type of positive ISH reaction in nuclei that appear normal by H&E staining was the only type of reaction reported by Cruz-Flores et al. (2022) in their PvSV-infected specimens. They described no circular inclusions in HP cytoplasmic vacuoles similar to those we found that were similar to the TEM image of a circular vacuolar inclusion containing PvPV virions in the publication by Liu et al. (2021). We are uncertain about the relationship between the ISH reactions in vacuolar inclusions versus those in nuclei, but we hope that TEM may provide some insight. In addition, it is possible that the positive ISH reactions might arise from endogenous viral elements (EVE) of WzSV8 within the genomic DNA of the respective RT-PCR positive specimens. For example, EVE of WzSV8 have been reported from *P. monodon* in Australia (Huerlimann et al., 2018).

In summary, our ISH assays in WzSV8 RT-PCR positive shrimp revealed that only the unusual lesions that occurred in E-cells were morphologically and locationally distinctive enough to be useful for rapid screening of H&E-stained tissues for matching lesions. These lesions consisted of circular, ISH-positive inclusions within cytoplasmic vacuoles of tubule epithelial cells in the E-cell region, and we could use the location and morphology of the lesions to search H&E-stained tissues for cells with similar lesions.

### WzSV8 lesions in H&E-stained tissues have some unique characteristics

H&E-stained tissue sections adjacent to the sections that had given positive ISH reactions were screened for the presence of lesions characterized by circular inclusions within cytoplasmic vacuoles of tubule epithelial cells in the E-cell region. Thus, we discovered cytoplasmic, deeply basophilic inclusions that were circular in section and were contained within a cytoplasmic vacuole. These are illustrated in **Figs. 7 and 8** in which tangential HP sections reveal HP tubules at their distal regions where B-cells, R-cells and F-cells are absent. These inclusions should not be confused with metaphase plates in plane section (**Fig. 9**) commonly seen in E-cells where there is a high rate of cell division. Although these deeply basophilic, circular inclusions of WzSV8 were dominant in the lesions, there were also variations in lesion appearance. For example, some lesions showed the basophilic inclusion associated with a nearby or attached, eosinophilic, circular inclusion, usually of smaller size (**Fig. 10**) (i.e., “double-inclusions”).

**Figure 7.**
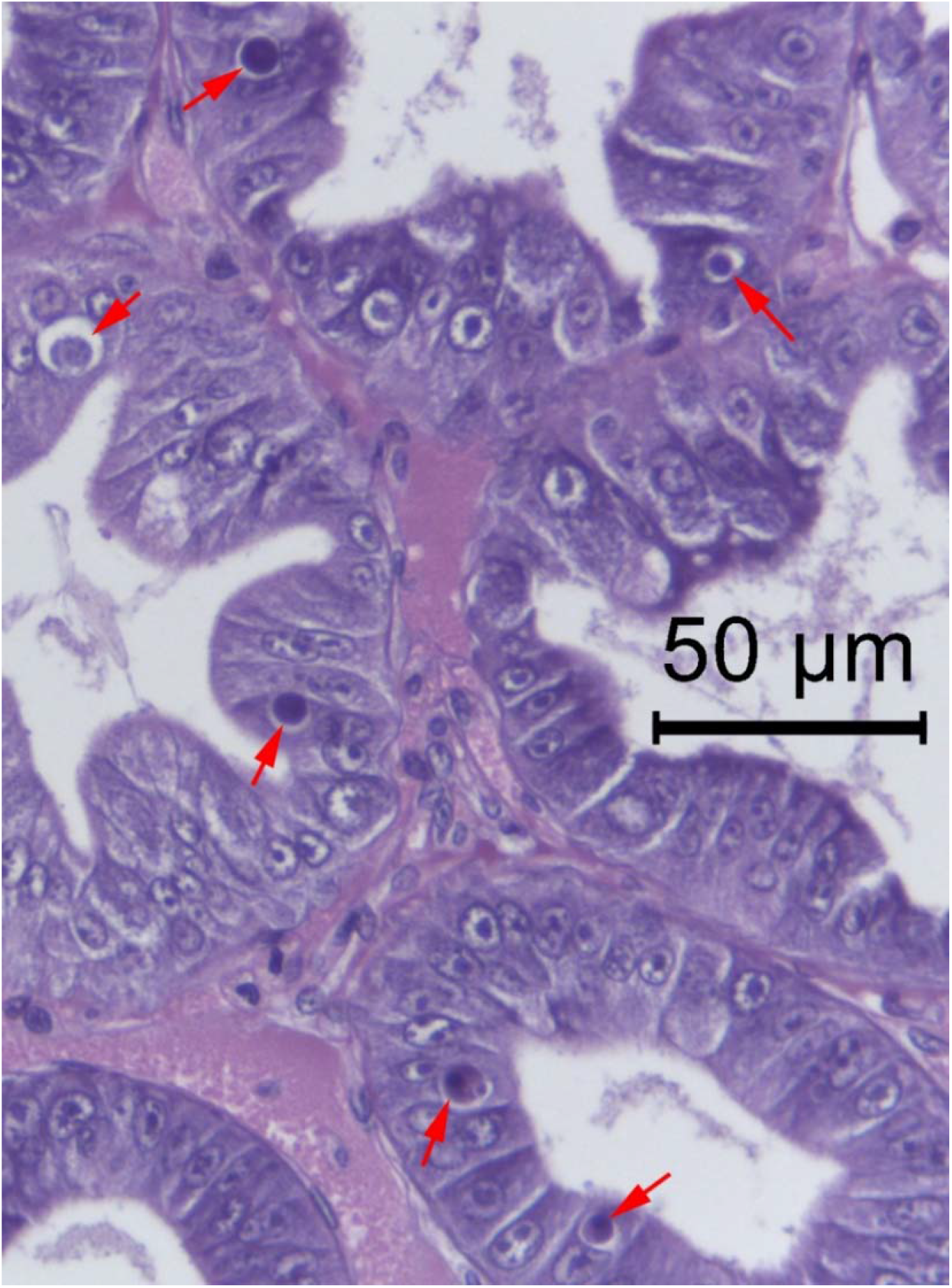
Photomicrograph taken using a 40x objective showing E-cells of the shrimp hepatopancreas containing WzSV8 cytoplasmic inclusions within vacuoles (red arrows). They have variable morphology and staining properties. The dominant ones are deeply basophilic and mostly perfectly circular. The lightly basophilic inclusion on the upper far left may be an early developmental stage, while the bottom two are more complex and show some eosinophilic staining in addition to a central, circular, densely basophilic inclusion.

**Figure 8.**
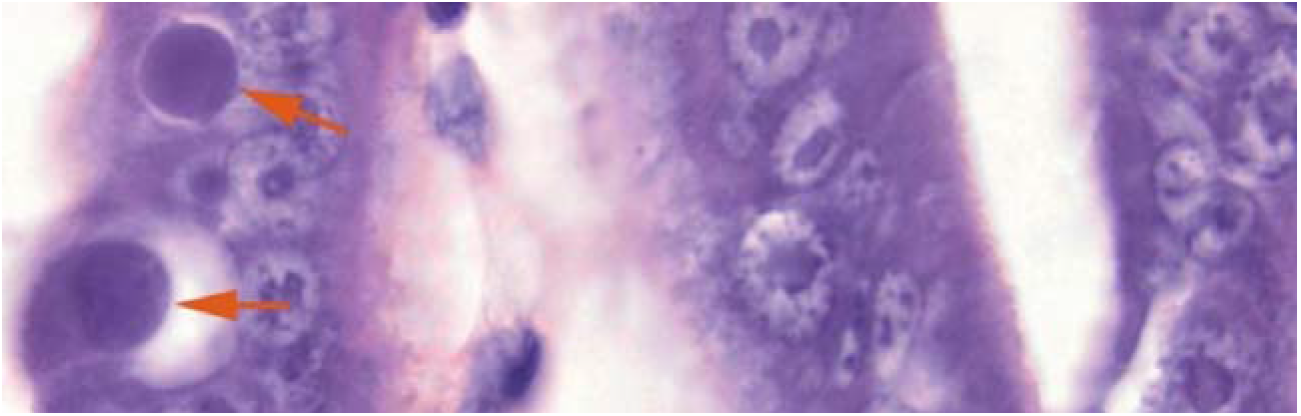
Example photomicrographs of the most common, circular, lightly to deeply basophilic, cytoplasmic inclusions of WzSV8 within vacuoles in E-cells.

**Figure 9.**
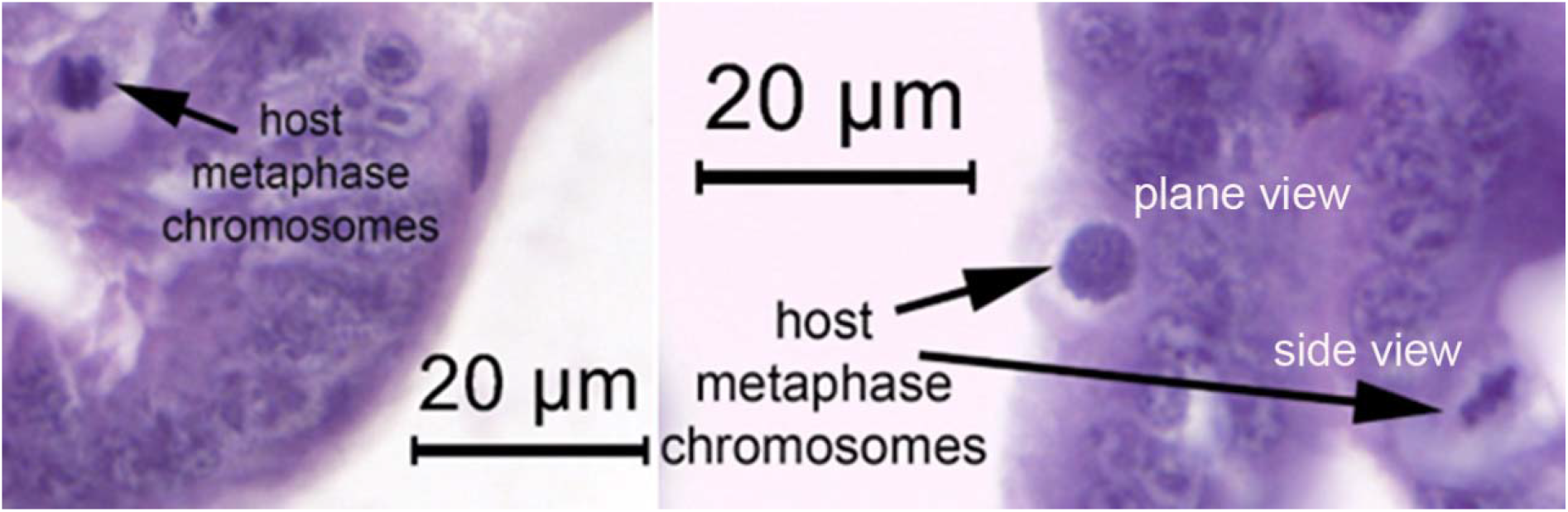
Example photomicrographs of metaphase chromosomes in E-cells that may sometimes resemble WzSV8 inclusions when the tissue section passes through the plane of the metaphase plate rather than the side. One must be careful not to confuse these with WzSV8 inclusions.

**Figure 10.**
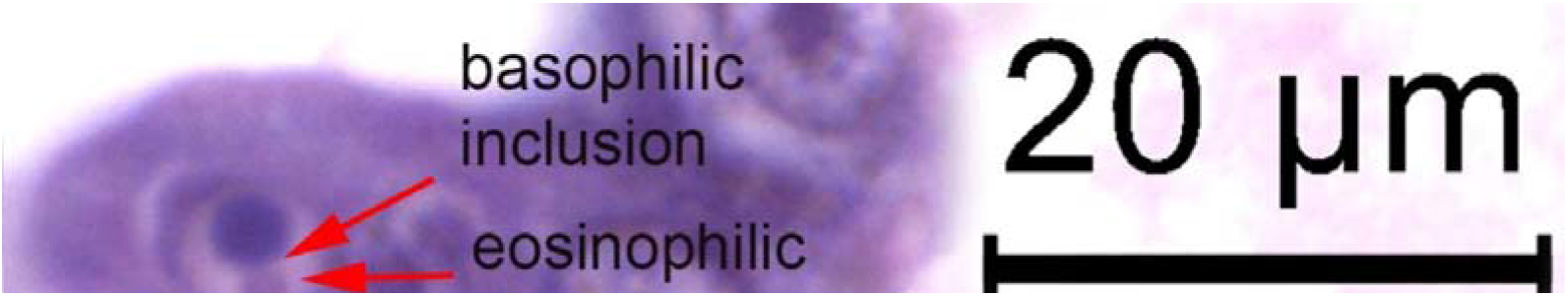
Variations in basophilic WzSV8 inclusions that are sometimes accompanied by usually smaller, circular, eosinophilic satellite inclusions. This is a highly distinctive combination, such that “densely basophilic, circular inclusions accompanied by a closely associated satellite, eosinophilic inclusion within an E-cell vacuole” may be considered pathognomonic for WzSV8.

Sometimes the lesions were also seen in differentiated HP tubule epithelial cells (confirmed by ISH), and an example photomicrograph of WzSV8 inclusions in R-cells is shown in **Fig. 11**. In addition, the double-inclusions in some lesions were sometimes separated by an unstained space from a surrounding basophilic to magenta colored “surround” of variable thickness that was itself surrounded by an unstained space. We are uncertain as to the steps in development of the “surround” but speculate that it may consist of the whole cell cytoplasm that has shrunken away (preparation artifact?) from adjacent cells. Hopefully, further TEM investigations will clarify this issue.

**Figure 11.**
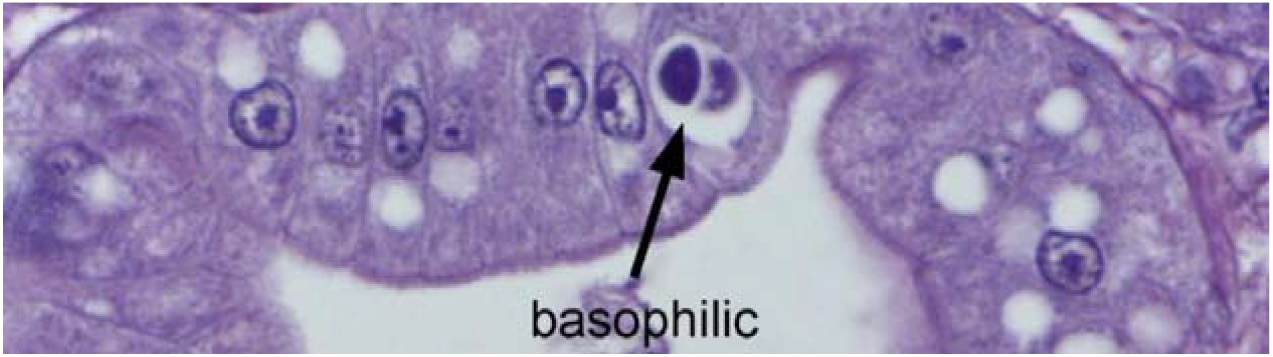
Photomicrograph of WzSV8 basophilic inclusions in differentiated cells (R-cells) of the shrimp hepatopancreas.

The double-inclusions consisting of a basophilic and eosinophilic partner may be common, but the eosinophilic partners are mostly small in comparison to the larger basophilic inclusion such that the probability of them appearing together in 4-micron tissue sections would be small. Thus, it is difficult to determine whether this unique pairing occurs regularly or is of low occurrence. More work is needed to determine the variation in elements that accompany the mostly circular ISH positive inclusions and how they develop. For example, it is likely from the TEM work by Liu et al. (2021) that the circular basophilic inclusion consists of WzSV8 virions, but it is not known whether the eosinophilic satellite originates from the virus itself, or from the host in response to the viral infection. Hopefully, TEM will help to clarify their nature and origin. However, this does not detract from the diagnostic value of their unique occurrence. For example, the Cowdry A type inclusion (an intranuclear, eosinophilic inclusion surrounded by an unstained space that separates it from the basophilic, maginated chromatin) is useful in identifying the shrimp virus IHHNV but is an artifact that arises from the use of acidic fixatives like Davidson’s fixative (Lightner 1996). Despite being an artifact, the Cowdry A type inclusion is considered to be a useful character for detecting or diagnosing IHHNV infections

Photomicrographs of semi-thin sections of WzSV8 inclusions in E-cells are also shown in **Fig. 12** stained with toluidine blue. The double-inclusions are so distinctive in their nature by both H&E staining and in semi-thin sections that their occurrence, together with their common location in E-cells may be considered pathognomonic for WzSV8 infection. We propose calling these unique double inclusions “Lightner double inclusions” (LDI) to honor recently deceased Prof. Donald V. Lightner to whom we are greatly indebted for his monumental contributions in the field of shrimp pathology.

**Figure 12.**
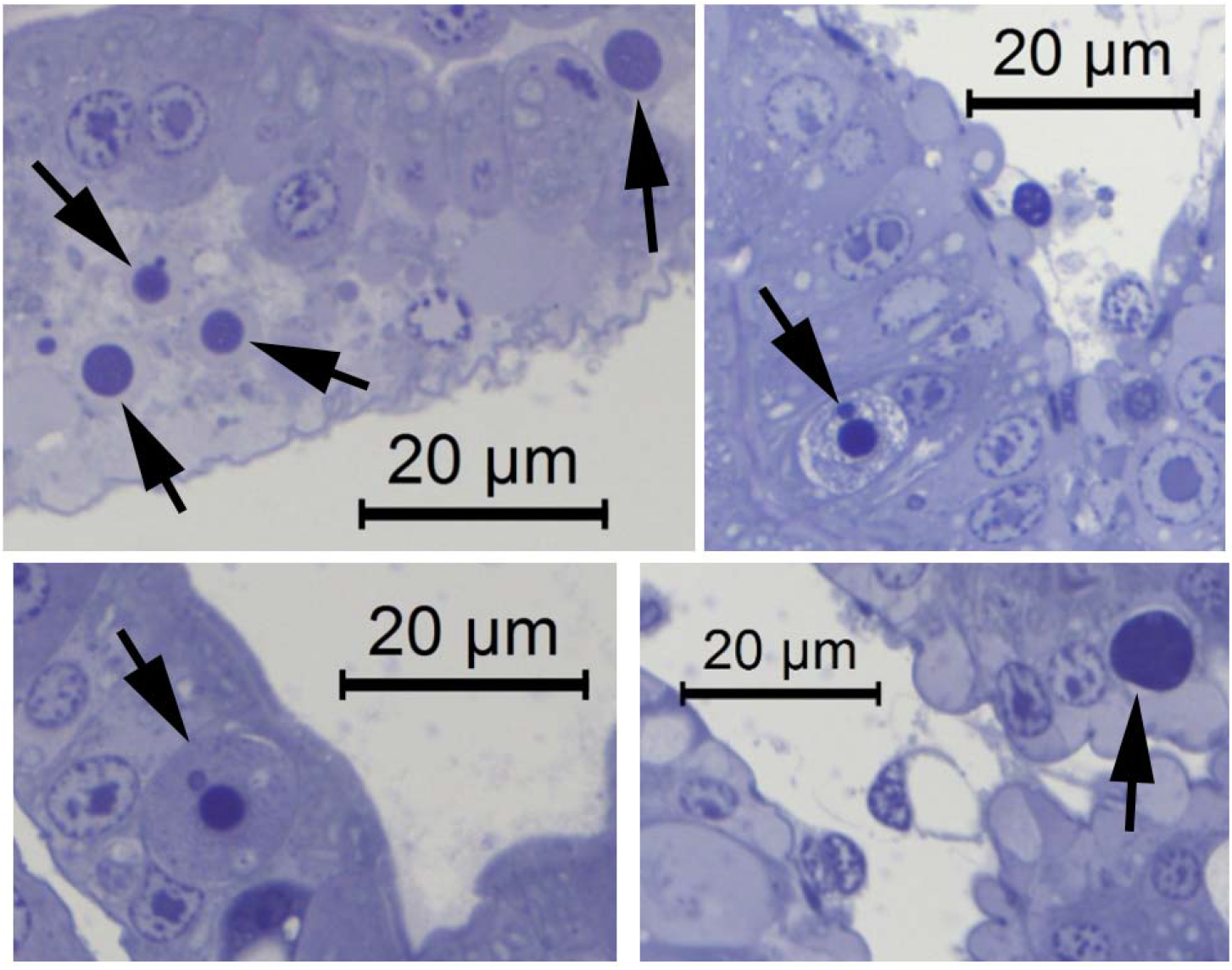
Photomicrographs of semi-thin sections of shrimp HP tissue showing variation in WzSV8 inclusions (dark blue and circular) in E-cells of the shrimp hepatopancreas. Some show a smaller, adjacent satellite inclusion or other vacuolar contents.

### Retrospective on WzSV8 inclusions

In retrospect, we had previously seen WzSV8 inclusions in both *P. monodon* and *P. vannamei* from several shrimp farming countries in Austral-Asia since at least 2008. They were described as of unknown origin in histological reports to clients. More recently we obtained samples of *P. vannamei* from the Americas that also showed these inclusions. We regarded them as “mystery inclusions” that we originally speculated might be developmental stages of the microsporidian *Enterocytozoon hepatopenaei* (EHP). However, the “mystery inclusions” were subsequently found to be negative for EHP using a specific ISH probe for EHP (unpublished). In addition, we had already discovered by histological analysis and ISH assays that EHP does not infect E-cells (Flegel, 2012; Chaijarasphong et al., 2020). Thus, the inclusions remained a mystery.

In most cases, occurrence of the “mystery inclusions” was not associated with disease. When they were present in diseased shrimp, they occurred together with other, known lethal pathogens (bacteria or viruses) that were deemed the cause of morbidity. Lacking a clear link to disease and being sporadic in occurrence, we regarded them somewhat as a curiosity that was not pursued due to other more urgent work.

The original publication describing WzSV8 did not include information related to signs of disease, histopathology or pathogenicity (Li et al. 2015). Nor did the publications on PvPV (Liu et al. 2021) and PvSV (Cruz-Flores et al., 2022). However, to confirm virulence, isolation of a new virus from moribund shrimp must be accompanied by histopathological analysis for the presence of other known pathogens together with results from challenge tests employing the new purified virus to show that it alone can produce the same disease as seen in the original diseased shrimp. To date, there is no published proof that WzSV8-related viruses have caused disease, and we have many histological samples from normal shrimp dating back over a decade that show the distinctive WzSV8 LDI. This suggests that WzSV8-like viurses may have had little impact on production of cultivated *P. monodon* and *P. vannamei*.

Despite historical evidence indicating lack of virulence, it is possible that many types of WzSV8 exist and that some may be lethal or may contribute to mortality in combination with one or more other pathogens or under some environmental conditions. It is also possible that a new, more virulent type is emerging. For example, yellow head virus (YHV) is known to occur in up to 8 types (Walker et al., 2021) but only one type is listed as reportable to the World Organization for Animal Health. This is what needs to be established as quickly as possible. Thus, it is important to mobilize researchers in all shrimp rearing countries to assess the distribution and potential virulence of WzSV8-like viurses. We hope that retrospective review of existing histological samples for LDI together with RT-PCR screening of fresh material will lead to rapid acquisition of new information and to the development of a more universal RT-PCR detection method. At the same time, it is important to keep in mind that a rapid rise in reports of WzSV8 occurrence will not be an indication of viral spread but simply the discovery of its current range. This virus has been around for at least a decade or more with no apparent causal link to disease.

## Acknowledgement

This research project is supported by Mahidol University (Fundamental Fund: Basic Research Fund: fiscal year 2022) (Grant no. BRF1-054/2565) and the NSRF via the Program Management Unit for Human Resources & Institutional Development, Research and Innovation (B05F640137). We also thank Agricultural Research Development Agency (ARDA), Thailand (PRP6505030760) to Kallaya Sritunyalucksana.

## Supplementary Figures for Wenzhou virus 8 (WzSV8) detection

**Supplementary Fig. S1.**
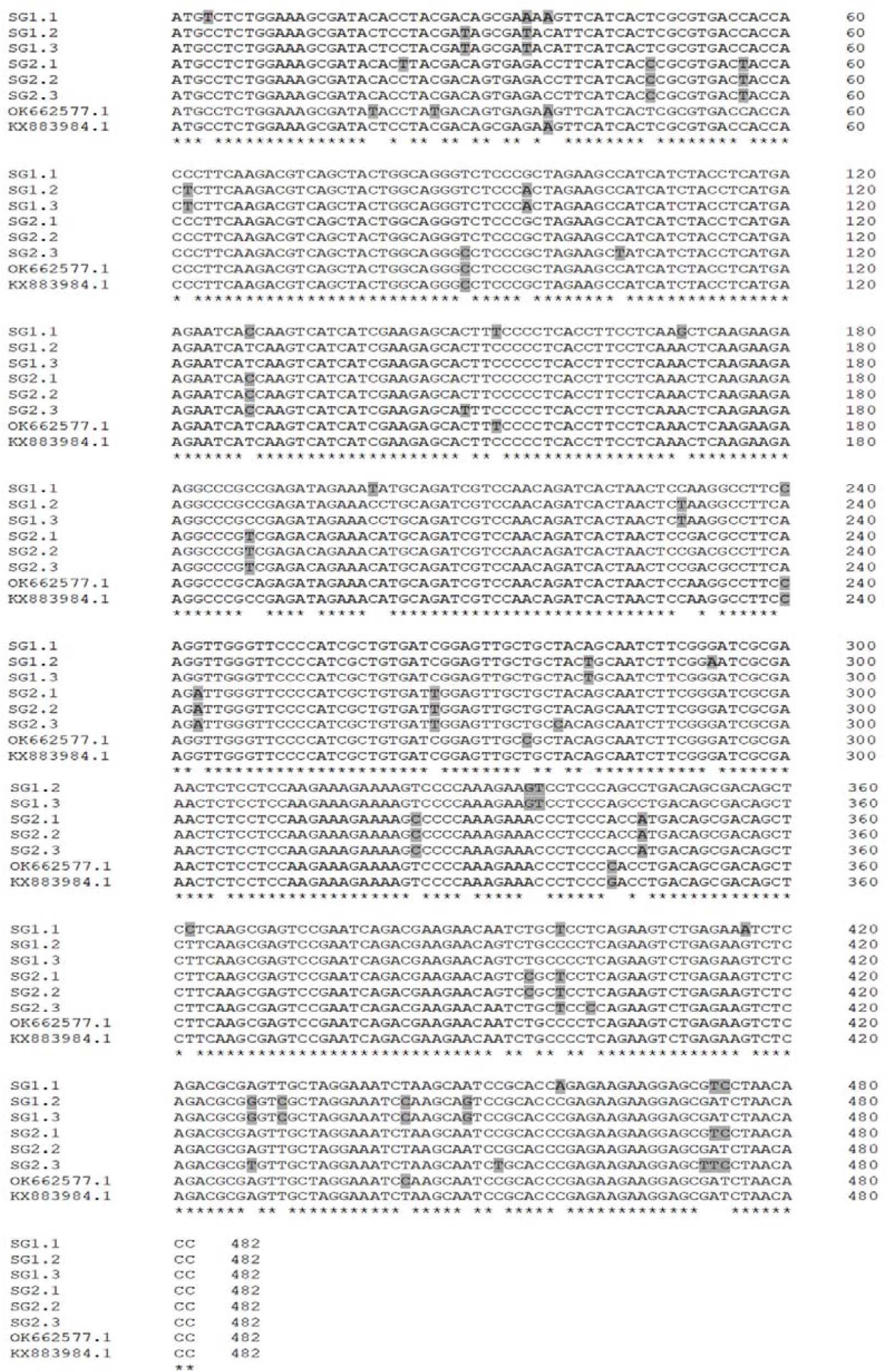
Clustal Omega alignment of the sequences of 482 bp-amplicon obtained using the method herein from specimens of Indo-Pacific (SG1) and America (SG2) origin with the two GenBank records for WzSV8 (KX883984.1) and *Pv*PV (OK662577.1) from China. The gaps in the line of asterisks indicate regions of difference among the 8 sequences, while the grey background nucleotides highlight nucleotide differences among the eight sequences. The % identities are shown in Table 2.

**Supplementary Figure S2.**
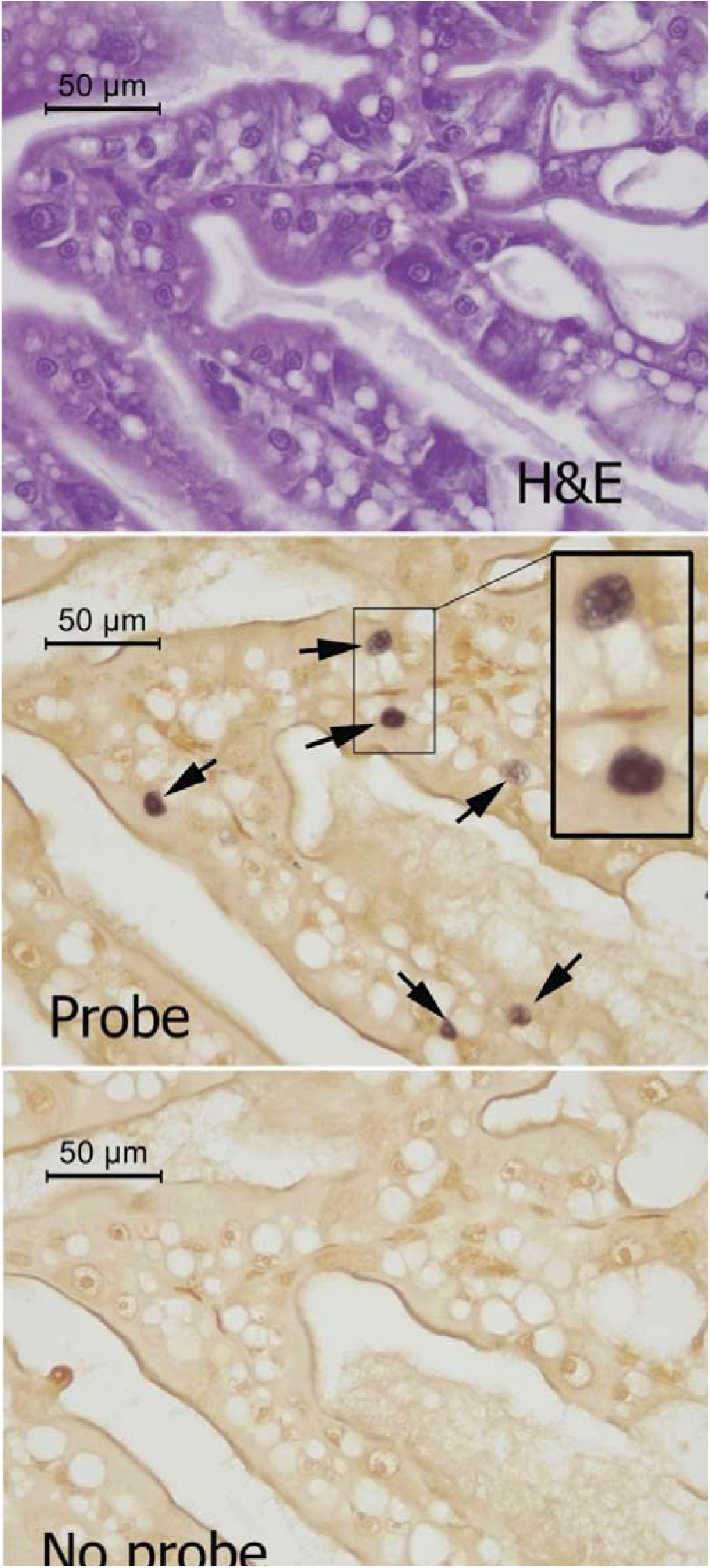
Photomicrographs of adjacent tissue sections from the medial region of the hepatopancreas (HP) of a shrimp specimen positive for WzSV8 by RT-PCR This shows *in situ* hybridization (ISH) signals (dark staining) in normal nuclei in normal tubule epithelial cells. The cells and nuclei have normal morphology in the hematoxylin and eosin (H&E) stained adjacent section, Thus, the presence of WzSV8 in specimens with such ISH positive nuclei would not be revealed by H&E staining, making H&E-stained sections of such cells of no use in diagnosis for the presence WzSV8 in such nuclei.

**Supplementary Figure S3.**
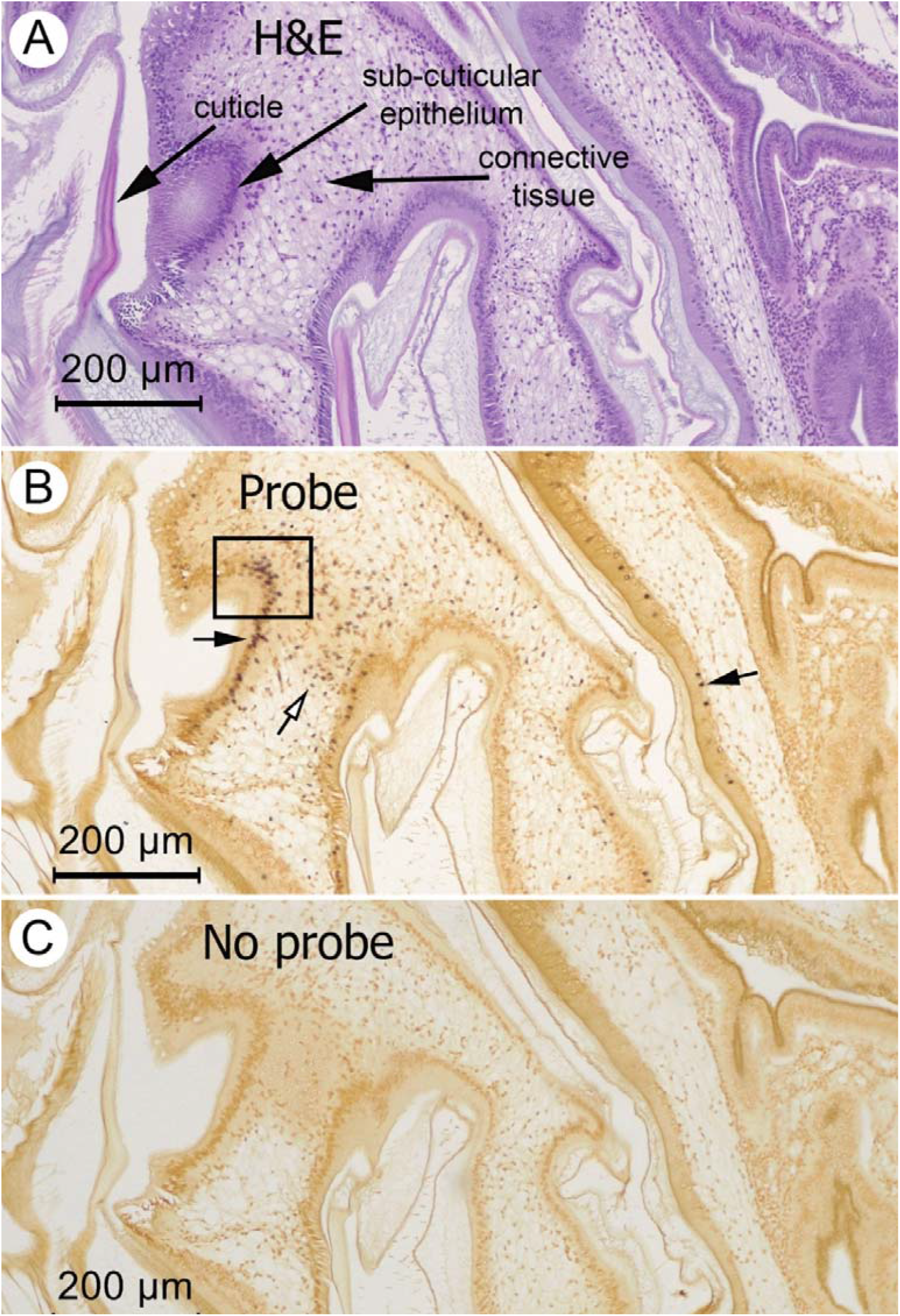
Photomicrographs of adjacent tissue sections from the region of the stomach of a shrimp specimen positive for WzSV8 by RT-PCR. These are low magnification photomicrographs of adjacent tissue sections stained with H&E and also tested for the presence of WzSV8 by ISH. A. H&E stained tissue section showing the stomach cuticle, sub-cuticular epithelium and connective tissue. B. Adjacent tissue section showing positive ISH reactions in nuclei of the sub-cuticular epithelium (black arrows) and underlying connective tissue (white arrow). The box indicates the area magnified in Supplementary Fig. 3. C. No probe control.

**Supplementary Figure S4.**
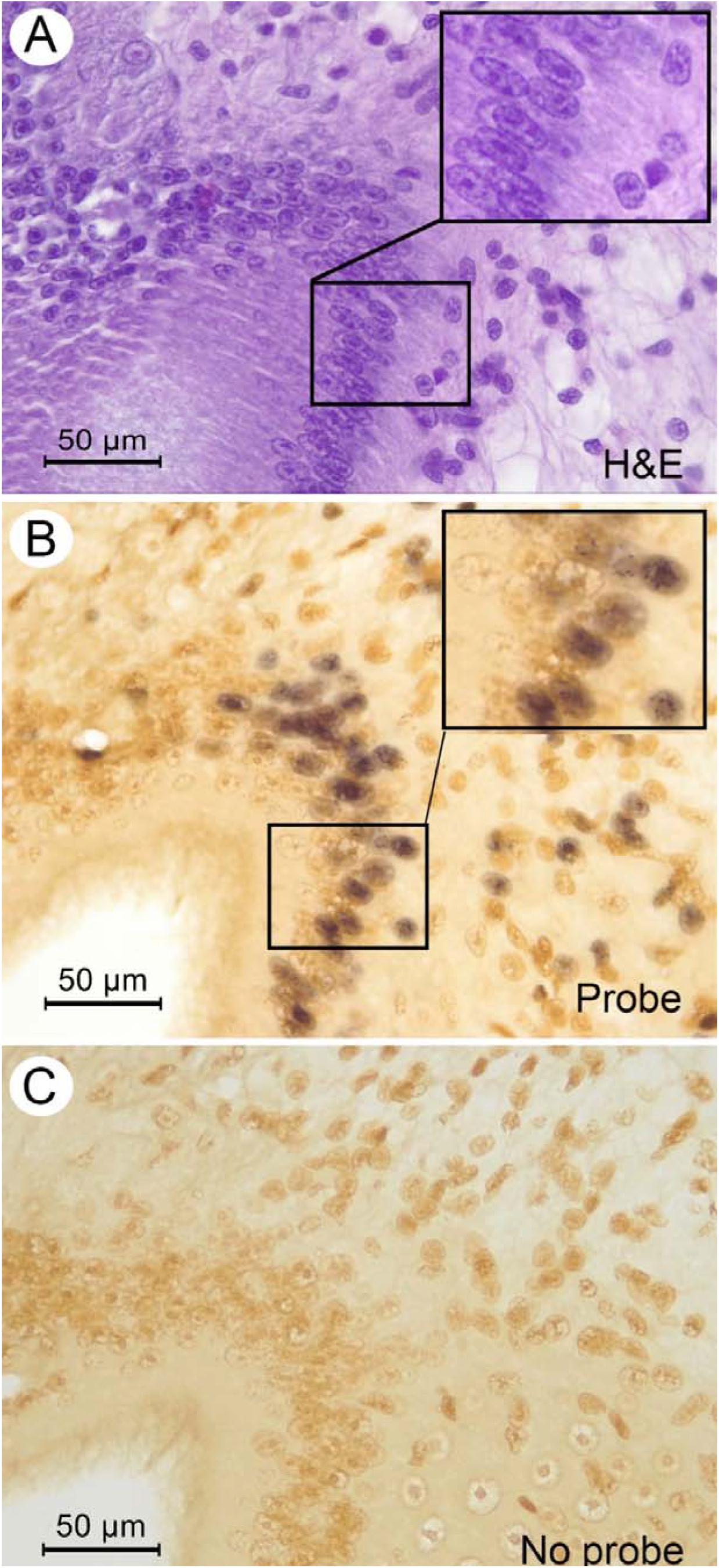
Photomicrographs of the stomach area at higher magnification indicated by the box in Fig. 2B. All show normal morphology of nuclei despite positive ISH reactions shown in B. A. H&E-stained tissue section. B. Adjacent tissue section showing positive ISH reactions in nuclei (dark staining), C. Adjacent tissue section for the no-probe negative control.

## REFERENCES

Bell, T.A., Lightner, D.V., 1988. A handbook of normal shrimp histology. World Aquaculture Society, Baton Rouge, LA.

Chaijarasphong, T., Munkongwongsiri, N., Stentiford, G.D., Aldama-Cano, D.J., Thansa, K., Flegel, T.W., Sritunyalucksana, K., Itsathitphaisarn, O., 2020. The shrimp microsporidian Enterocytozoon hepatopenaei (EHP): Biology, pathology, diagnostics and control. J. Invertebr. Pathol., 107458.

Cruz-Flores, R., Andrade, T.P., Mai, H.N., Alenton, R.R.R., Dhar, A.K., 2022. Identification of a novel solinvivirus with nuclear localization associated with mass mortalities in cultured whiteleg shrimp (Penaeus vannamei). Viruses. 14, 2220.

Flegel, T.W., 2012. Historic emergence, impact and current status of shrimp pathogens in Asia. J. Invertebr. Pathol. 110, 166–173.

Huerlimann, R., Wade, N.M., Gordon, L., Montenegro, J.D., Goodall, J., McWilliam, S., Tinning, M., Siemering, K., Giardina, E., Donovan, D., 2018. De novo assembly, characterization, functional annotation and expression patterns of the black tiger shrimp (Penaeus monodon) transcriptome. Sci. Rep. 8, 1–14.

Li, C.-X., Shi, M., Tian, J.-H., Lin, X.-D., Kang, Y.-J., Chen, L.-J., Qin, X.-C., Xu, J., Holmes, E.C., Zhang, Y.-Z., 2015. Unprecedented genomic diversity of RNA viruses in arthropods reveals the ancestry of negative-sense RNA viruses. eLife. 4, e05378.

Lightner, D.V., 1996. A handbook of pathology and diagnostic procedures for diseases of penaeid shrimp. World Aquaculture Society, Baton Rouge, LA.

Liu, S., Xu, T., Wang, C., Jia, T., Zhang, Q., 2021. A novel picornavirus discovered in white leg shrimp Penaeus vannamei. Viruses. 13, 2381.

Pénzes, J.J., Söderlund-Venermo, M., Canuti, M., Eis-Hübinger, A.M., Hughes, J., Cotmore, S.F., Harrach, B., 2020. Reorganizing the family Parvoviridae: a revised taxonomy independent of the canonical approach based on host association. Arch. Virol., 1–14.

Srisala, J., Sanguanrut, P., Thaiue, D., Laiphrom, S., Siriwattano, J., Khudet, J., Powtongsook, S., Flegel, T.W., Sritunyalucksana, K. 2021. Infectious myonecrosis virus (IMNV) and Decapod iridescent virus 1 (DIV1) detected in captured, wild Penaeus monodon. Aquaculture. 545, 737262.

Sriurairatana, S., Boonyawiwat, V., Gangnonngiw, W., Laosutthipong, C., Hiranchan, J., Flegel, T.W., 2014. White feces syndrome of shrimp arises from transformation, sloughing and aggregation of hepatopancreatic microvilli into vermiform bodies superfificially resembling gregarines. PLoS One. 9 (6), e99170.

Walker, Peter J, Jeff A Cowley, Xuan Dong, Jie Huang, Nick Moody, John Ziebuhr, and ICTV Report Consortium. 2021. ICTV Virus Taxonomy Profile: Roniviridae. J Gen Virol, 102.

